# The structure and function of neural connectomes are shaped by a small number of design principles

**DOI:** 10.1101/2023.03.15.532611

**Authors:** Adam Haber, Adrian A. Wanner, Rainer W. Friedrich, Elad Schneidman

## Abstract

The map of synaptic connectivity among neurons in the brain shapes the computations that neural circuits may perform. Inferring the design principles of neural connectomes is, therefore, fundamental for understanding brain development and architecture, neural computations, learning, and behavior. Here, we learn probabilistic generative models for the connectomes of the olfactory bulb of zebrafish, part of the mouse visual cortex, and of *C. elegans*. We show that, in all cases, models that rely on a surprisingly small number of simple biological and physical features are highly accurate in replicating a wide range of properties of the measured circuits. Specifically, they accurately predict the existence of individual synapses and their strength, distributions of synaptic indegree and outdegree of the neurons, frequency of sub-network motifs, and more. Furthermore, we simulate synthetic circuits generated by our model for the olfactory bulb of zebrafish and show that they replicate the computation that the real circuit performs in response to olfactory cues. Finally, we show that specific failures of our models reflect missing design features that we uncover by adding latent features to the model. Thus, our results reflect surprisingly simple design principles of real connectomes in three different systems and species, and offer a novel general computational framework for analyzing connectomes and linking structure and function in neural circuits.

## Introduction

The ability to reconstruct the detailed connectivity maps between neurons at unprecedented scale and resolution (1–5) opens the door for the quantitative analysis of the organization of real neural networks, the inference of their structural design principles, and the uncovering of the relations between structure and function in neural circuits (6–8). Analyses of the detailed architecture of brain networks have shown that synaptic connectivity is structured at different scales: from the enrichment of reciprocal connections (9) and network motifs (10–12), through the structured organization of the retina (13), to cortical columns (14–16), and other nervous systems and brain areas (1,17–19). A wide range of genetic (20–23), morphological (24, 25), biochemical (26), and economical (27) mechanisms have been implied or shown to play a significant role in shaping the structure of these networks, but the interplay between these mechanisms is still not well understood. Moreover, it remains unclear to what extent the large-scale organization of brain networks can be explained by simple and local connectivity rules. Complementary to this “bottom-up” approach, network theory tools (28, 29) have been used to study the design of networks of neurons (30, 31), yielding models able to reproduce global properties, such as “small-worldness” (32) and modular organizations (33). However, it is unclear how these properties would emerge from local synaptic formation mechanisms, such as the ones described above. Here, we merge these two viewpoints of network analysis and design by learning generative models of large connectomes whose basic building blocks correspond to simple and natural biological structural features and physical constraints.

Inferring and understanding the the design principles of networks and their organization require a modeling formalism that would allow us to bring together local rules, global network structures, and the inherent stochastic nature of individual networks. This is because any given connectome is the result of genetic instructions, developmental processes, and learning throughout the lifetime of the organism – which are all noisy biophysical processes that also depend on other external conditions. It is clear then that the detailed connectivity of the same areas across animals would be probabilistic in nature, and no two circuits would have exactly the same neurons or connections (except, maybe, for specialized circuits with “identified” neurons). So, to describe and evaluate connectomes, we must rely on statistical models over the possible connectivity maps. Such maps of connections between neurons are naturally described as a directed graph or a matrix *G*, where each entry *G_ij_* reflects the synaptic connections from neuron *i* to neuron *j*. A statistical model of connectomes would then be a probability distribution over connectivity matrices, *G*, that depends on a set of parameters *θ.*

Learning such accurate models for connectomes, *P_θ_*(*G*), would allow us to (1) evaluate the likelihood of a particular connectivity map, (2) generate synthetic connectomes that resemble the observed ones, by sampling from the model, (3) quantify the expected values of different features of the connectome and their variability, and (4) explore the functional design of connectomes, by simulating the neural dynamics of the sampled networks. From a computational perspective, we would seek models that recapitulate the observed structure as accurately as possible, and identify the minimal set of features that would be needed. From a biological perspective, we would like to find models that rely on biologically-plausible or “understandable” features. Learning such statistical models is a challenging computational task,to which should be added in this study the hurdle of having only a few examples of reconstructed connectomes. Consequently, we aim to learn models based on the observed values of a small number of simple and biologically-plausible features or statistics of the connectome, as summarized in Fig. 1, and evaluate these models using a range of statistical and structural measures that we did not use for learning the models – at the level of individual synapses, individual neurons, small sub-circuits, and the circuit as a whole. We then ask how well these models predict the function of the circuit, what they might reveal about the design principles that we are still missing, and how we might find them.

**Fig. 1.**
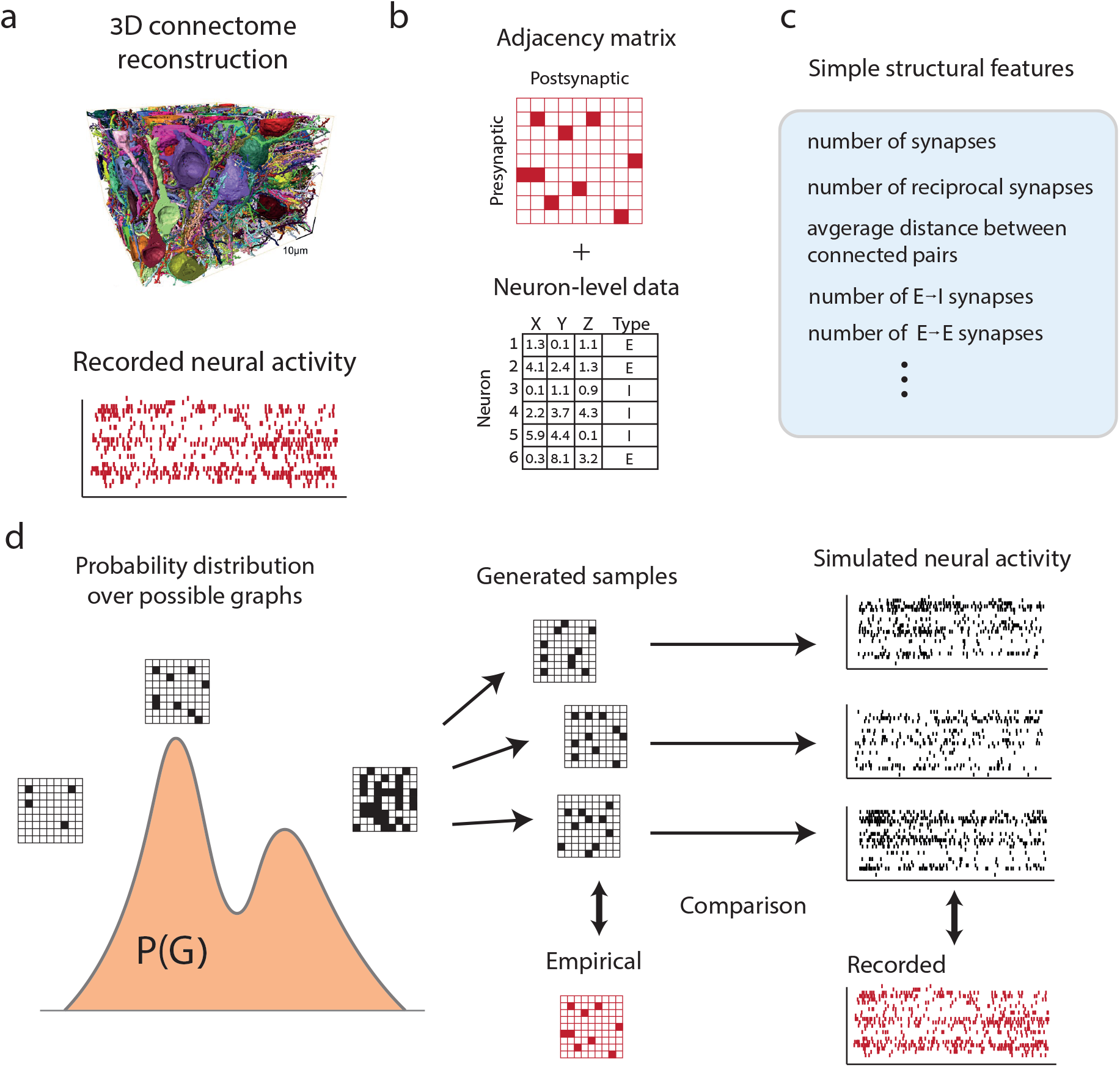
Learning and evaluating feature-based generative models for neural connectomes. **(a)** A complete 3D reconstruction of a neural tissue containing *N* neurons (adapted with permission from (3)). **(b)** The network of synaptic connections is represented as a *N* × *N* binary connectivity matrix, augmented by additional neuron-level data, such as the spatial positions of individual cells, cell types, etc. **(c)** A set of simple structural features is computed from the connectivity matrix and the neuron-level data. **(d)** The chosen features are used to construct the maximum entropy probability distribution over the space of graphs, which reproduces the observed values of the features in the data, but is otherwise maximally random. Generated connectomes are sampled from the distribution and compared against the empirical connectivity, and, where available, simulations of activity of the sampled connectomes are compared with the recorded neural activity of the real circuit.

## Results

### Learning accurate generative models of connectomes based on simple biological and physical features

We first consider the connectome of part of the olfactory bulb (OB) of a zebrafish larva, taken from (34). This reconstructed network comprises more than 40% of the entire OB, containing 208 excitatory mitral cells (MCs) and 238 inhibitory interneurons (INs) that are connected by 9919 synapses (out of 198,470 possible ones), and the activity of its principal neurons has been measured (as we will see below). We start by using the unweighted version of the connectome (Fig. 2a; see Methods), as the estimation of synaptic strengths from structural data is sometimes partial or noisy (35). The connectome in this case corresponds to a binary matrix, *G*, where *G_ij_* = 1 designates the existence of a synapse or multiple synapses from neuron *i* to neuron *j.*

**Fig. 2.**
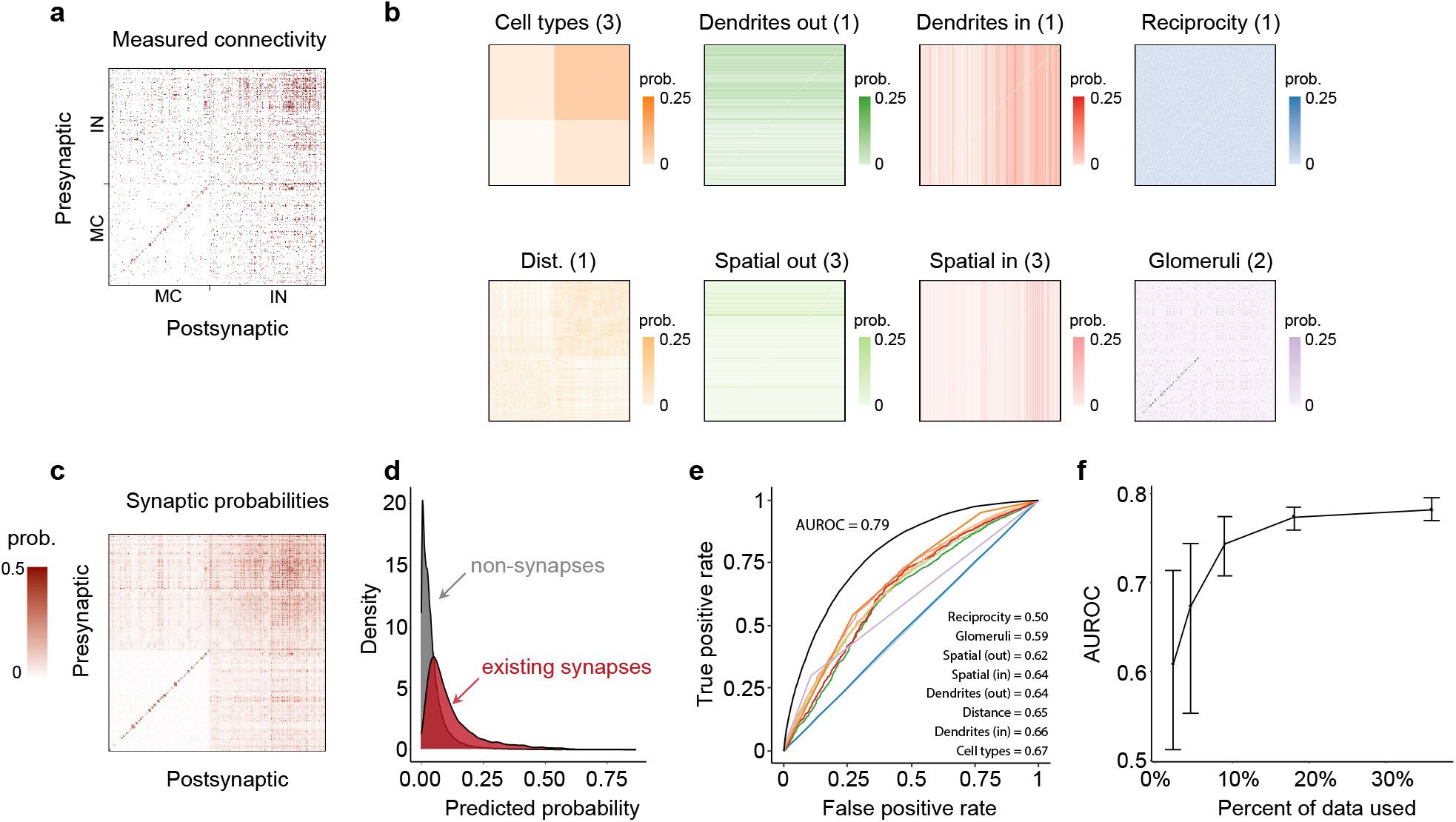
Accuracy of models based on different features in predicting individual synapses in the connectome of the olfactory bulb of zebrafish. **(a)** Synaptic connectivity map between 208 excitatory mitral cells (MCs) and 238 inhibitory interneurons (INs) from the olfactory bulb of the Zebrafish larva (34). **(b)** The synaptic probabilities predicted by the eight different tested models of the OB connectome, each defined using a different biological feature-set. Same ordering of the cells as in (a); numbers in parentheses correspond to the number of features in each feature-set **(c)** Synaptic probabilities predicted by the model that combines all eight feature-sets. See Supplementary Fig. S2 for individual samples from the models. **(d)** Distributions of synaptic probabilities assigned by the full model for empirically observed synapses (red) versus pairs of neurons that were not connected in the data (gray). **(e)** ROC curves for all single-feature models (same colors as in (b)), as well as the model that combines all the features (black). **(f)** Cross-validated AUROC of the full model as a function of the percentage of data used for training.

To identify the design principles of the OB connectome, we asked how well would models that rely on simple structural and biophysical features successfully replicate the observed connectivity. The simplest connectome model is one that relies solely on one structural feature – the total number of synapses in the network – and would adjust the overall sparsity of synaptic connections to match the observed one (4.9%). This is the well-known Erdős-Reńyi (ER) random graph model (28), which assigns the same probability to all potential synapses between neurons in the network. Because of its inherent homogeneity, this model gives a probability map for synaptic connections (i.e., a matrix containing the probabilities of synaptic connections between all pairs of neurons) that does not show any structure, (Supplementary Fig. S1), and is clearly a poor model of the real connectome. We then learn eight models for the connectome, that in addition to retaining the total number of synapses that were observed in the data, each model relies on a different set of features, and the model parameters are learned so that the expected feature values are consistent with their measured ones. Each of the eight feature-sets has a clear biological or physical interpretation: (1) cell-type-specific connectivity, (2) distance-dependent connectivity between neurons, (3) reciprocity of connections between pairs of neurons, (4) the dependence of incoming synapses on the location of the post-synaptic neuron, (5) the dependence of outgoing synapses on the spatial locations of the presynaptic neuron, (6) preferential attachment between Glomeruli, (7) the effect of dendritic tree sizes on the number of incoming synapses, and (8) the effect of dendritic tree sizes on the number of outgoing synapses (see Methods). Thus, for example, the model that relies on cell-type specific connectivity has 4 specific features – corresponding to the 2 x 2 values of the probability of a neuron of one type to have a synapse with a neuron of another type (as the neurons in this data set were identified as either MCs or INs). To assess the effect of each feature-set, we learn in each case the most random or least structured probabilistic model over networks, *P_θ_*(*G*), which matches the observed values of the corresponding features. Thus, given a set of features {*f_μ_*(*G*)}, we find the maximum entropy distribution over networks such that the average values of these features over the model and over the given connectome agree, namely, 〈*f_μ_*(*G*)〈*P* = 〈*f_μ_*(*G*)〉_*data*_. This model is given by

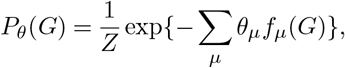

where the *θ_μ_*s are found numerically and *Z* is a normalization term or partition function (see Methods). These models, also known as Exponential Random Graph Models (36), not only allow us to compute the likelihood of any given connectome, but also serve as generative models that can be used to sample connectomes that are consistent with their respective features. Importantly, because these are maximum entropy models (37), the solution is unique and assumes no other structure beyond the measured features. These models, therefore, capture the full predictive power of their respective set of features. Fig. 2b shows the probability maps over the full connectivity matrix of synapses for each of the these models. As is evident, they each capture some aspects of the connectivity matrix, but none of them is particularly accurate on its own.

The generality of this mathematical framework means that we can naturally extend it to learn a model for the connectome that relies on any combination of the different featuresets simultaneously (as in (38)). Fig. 2c shows the synaptic probability map of this model and its similarity to the data (shown in Fig. 2a). Importantly, while predictions of the model may reflect high variance or uncertainty regarding individual synapses, existing synapses were assigned higher probabilities, despite the fact that the model was only trained on the aggregate summary statistics of the connectivity matrix (Fig. 2d).

We next quantify the performance of the different models by their ability to predict individual synapses, as well as the connectivity profiles of individual neurons, sub-circuit properties, and the likelihood of the connectivity map of the whole circuit. We first asked how accurately they predict the existence of individual synapses. This was assessed using the Receiver-Operating Characteristic (ROC) curve, which shows the rate of correctly predicting the existence of a synaptic connection vs. the rate of incorrectly predicting one, for different threshold classification values of the model (Fig. 2e, Methods). Critically, to avoid over-fitting, we trained the models on the connectivity matrix between only half of the neurons (randomly chosen), and used the other half as test data on which we assessed the accuracy of each model. Notably, these models proved to be highly robust, as evidenced by the observation that learning a model using only ~ 10% of the neurons sufficed to predict the connectivity of the rest of the network at a similar level (Fig. 2f, Methods). We summarize the predictions of each model by the mean area under the ROC curve (AUROC), averaged across different train and test splits of the data – which quantifies the ability of models to assign higher probabilities to existing synapses compared to non-existing ones (Methods). The models that rely on a single set of features have a limited predictive power,ranging from AUROC values of 0.5 (no predictive power) to 0.67. For comparison, the AUROC values obtained after training and testing the models on the whole connectome were effectively the same for all eight models (Supplementary Fig. S3).

But the ROC curve of the model that combined all the features, shown in Fig. 2e-f, gives an AUROC value of 0.79. The high accuracy of this model raises two obvious questions: First, do we need all the features that we used for the model to achieve this level of accuracy? Second, what features are we still missing?

### Redundancy among structural features implies that a small set of them shapes the connectome

The different feature-sets that we explored are related to one another, as can be gleaned, for example, from the spatial locations of different types of neurons (Fig. 3a). This redundancy is clearly reflected by the ability of a model based on one feature-set to predict the observed values of the other feature-sets, shown in Supplementary Fig. S4. Fig. 3b further shows the pairwise correlation between the predicted connectivity matrices of all eight tested models – demonstrating that the probabilities assigned to the synaptic connections by the different models are often correlated.

**Fig. 3.**
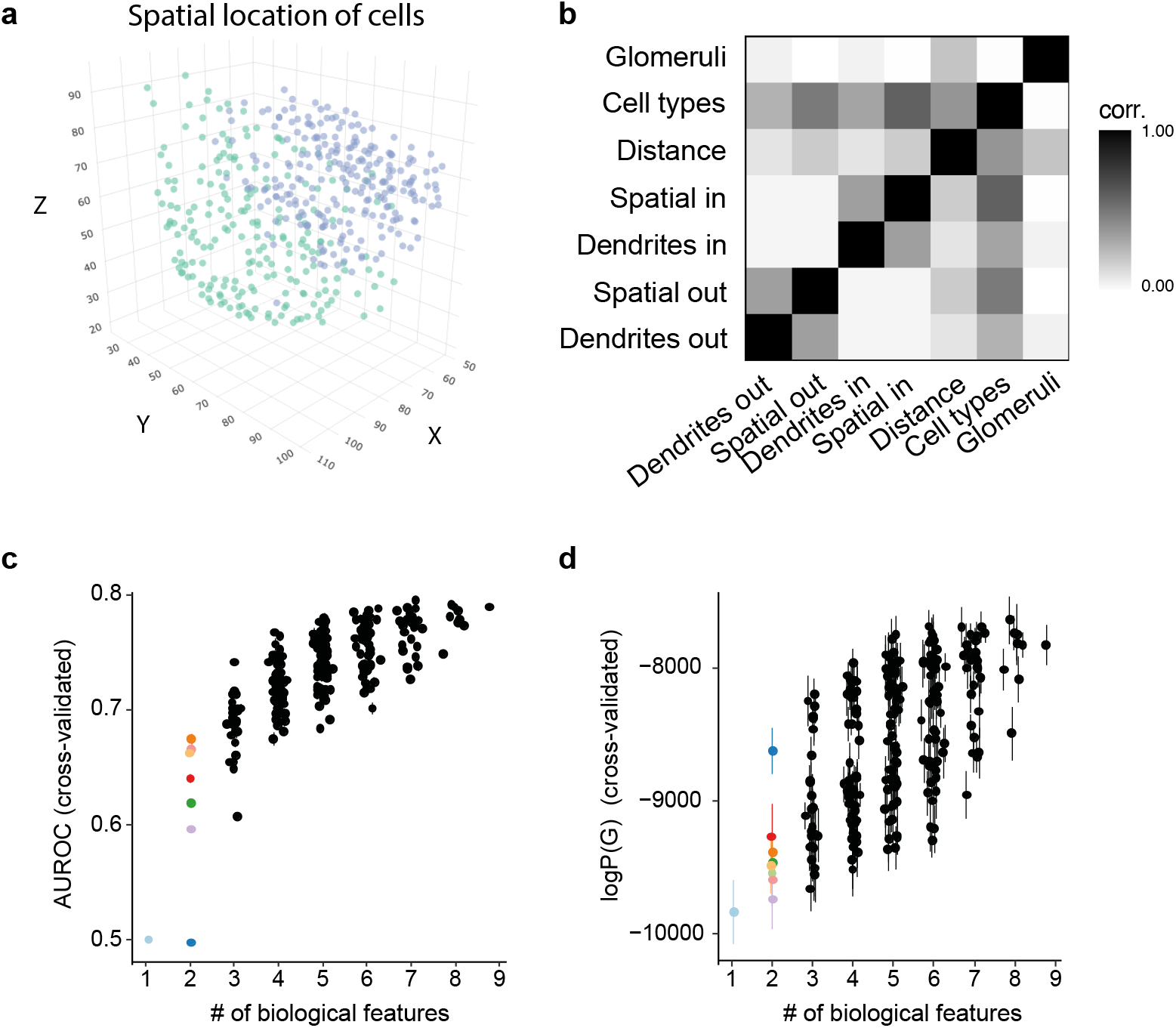
Redundancy among features imply that a small subset of them suffices to give highly accurate models of the OB connectome. **(a)** Spatial positions of cells in the OB connectome. Blue - INs; green - MCs. **(b)** Pearson correlation between the synaptic probability matrices of the different single feature-set models, computed between “flattened” matrices as vectors. The number-of-synapses model and the reciprocity model predict the same synaptic probability for all pairs and were, therefore, excluded from the analysis. **(c)** Cross-validated AUROC of all the 256 models in the ensemble. Colored dots represent the original single feature-set models (same colors as in Fig. 2b). **(d)** Same as in (c) but for log-likelihood scores.

Given this redundancy between features, we assess directly their necessity and sufficiency. This entailed comparing a large ensemble of models that are based on all the possible combinations of the different feature-sets, totalling 2^8^ = 256 different generative models. We find that while, in general, a greater number of features resulted in a higher accuracy of predicting synaptic connections, measured by their AUROC values (Fig. 3c), different combinations of features had very different values.

To go beyond predicting individual synapses, we also examine the performance of all 256 models on the full connectivity matrix. Using the log-likelihood values over the empirical connectome, log *P_θ_*(*G*), we find that also at the level of the whole connectome, adding features improves the likelihood values (Fig. 3d). Again, we emphasize that we trained the models on half of the connectome and predicted the connections in the other half (see SI for the performance on all the training data). For both measures, the “diminishing return” of adding features reflects the redundancy between features in shaping the connectome. Over the different models, likelihood values and AUROC values are highly correlated (Supplementary Fig. S5), but some of the outliers show the importance of individual features. For example, while the reciprocity does not contribute much to the AUROC score, it is critical for the likelihood of the model of the OB – reflecting the highly symmetrical nature of the connectivity.

We identify among all the feature combinations a model whose likelihood value is in the top 10% of all the models and is less than 5% away from the one that relies on all the features in terms of AUROC. This model is based on 5 of the feature sets: reciprocity, cell types, preferential attachment between Glomeruli, the effect of dendritic tree sizes on the number of incoming synapses, and the effect of spatial location on the number of outgoing synapses. We proceed with this particular set of features to explore other properties of the connectomes that this model generates, and ask would compact models of connectomes similarly apply to other species and neural systems.

### A small number of features accurately predict synaptic connectivity in different species

We perform a similar analysis on the reconstructed connectivity maps from two other species: (1) 334 L2/3 excitatory neurons from the mouse visual cortex (39). (2) chemical synapses of the neurons in the hermaphrodite *C. elegans* (40) (Methods).

As in the zebrafish analysis, we fit an ensemble of models that rely on all the combinations of the measured features: The mouse connectome had 1734 synapses, and the measured features were the spatial locations of the neurons and the distances between them, but no additional information on cell-types (Fig. 4a, green). For *C. elegans*, we used the connectome of 279 neurons (all non-pharyngal neurons), 3520 synapses, 15 cell-types, the 3D position of each neuron (Fig. 4a, blue) and its time of birth. For the cortical connectome, we did not use reciprocity or the dependence of incoming synapses on the spatial locations of neurons.

**Fig. 4.**
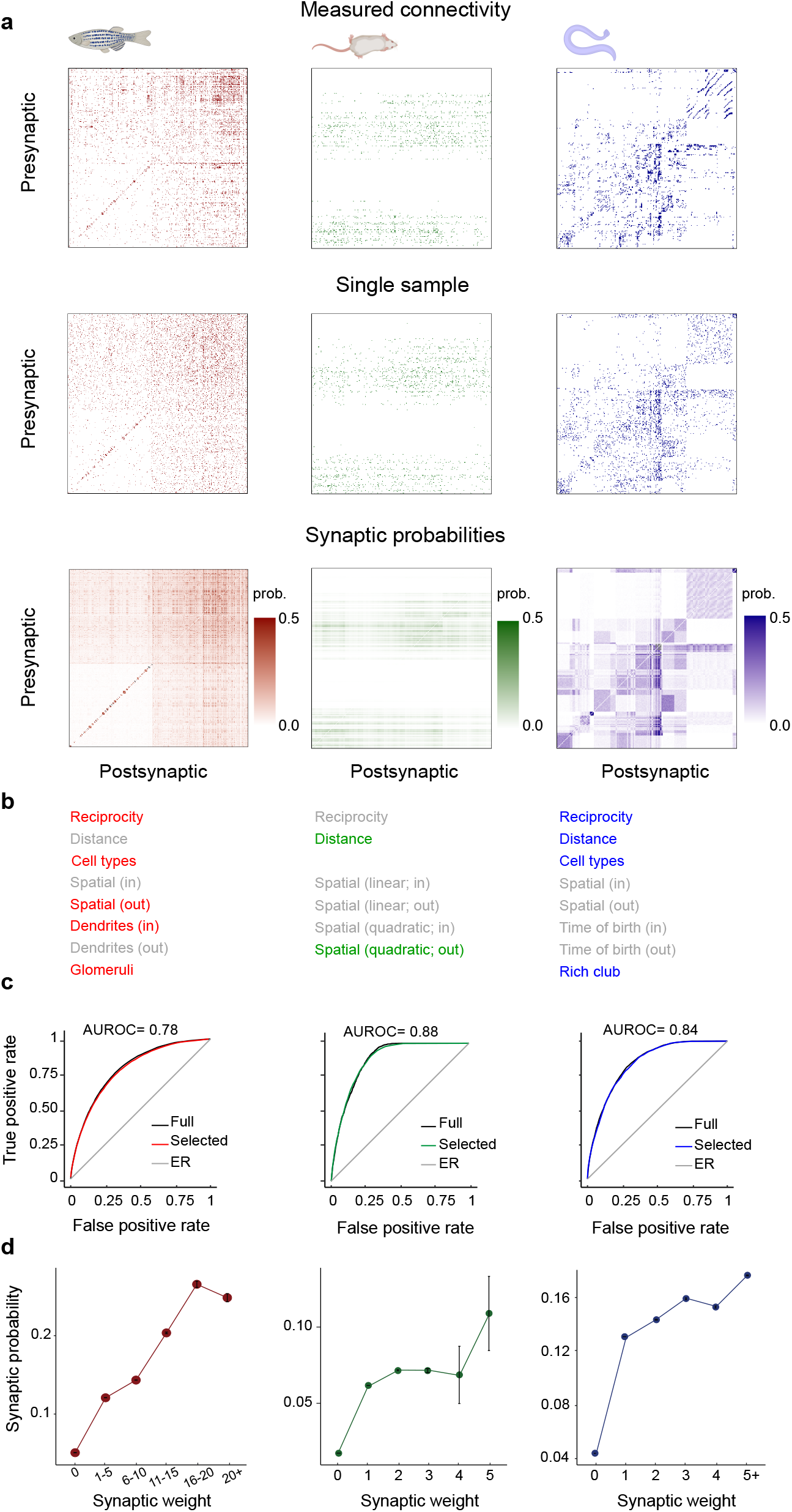
Models based on small number of feature-sets accurately predict synaptic connectivity and strength in the connectomes of zebrafish, mouse, and *C. elegans.* **(a)** The empirical connectivity matrix (top), a single sample from the selected model (middle), and the matrix of synaptic probabilities (bottom) for the fish (red), mouse (green) and worm (blue) connectomes. **(b)** The structural features we considered for each dataset. Colored features correspond to the features of the selected model for each dataset. **(c)** ROC curves of the respective selected model (colored) and the model that uses all the respective features (black) for each of the three connectomes. **(d)** The predicted synaptic probabilities correlate with the empirical synaptic weights (binned). Error bars represent one standard error of the mean, computed for all synapses within each bin.

In both the mouse and the worm, as in the fish, models based on small sets of features proved to be highly accurate: In the case of the cortical circuit, a model based on distance-dependent connectivity and a quadratic dependence of outgoing synapses on the spatial locations of the cells achieved an AUROC score of 0.88 (on cross-validated data). As for the *C. elegans* connectome, a model that relied on cell-type-specific connectivity, reciprocity, distance between cells, and the effect of membership in the C. *elegans* “rich-club” neurons (41) was enough to achieve an AUROC score of 0.84. Again, the redundancy between features allowed us to pick for each of the connectomes a compact combination of features that were in the top 10% of the models in terms of both their log-likelihood and AUROC value (normalized with respect to the model with the maximum AUROC) (Fig. 4c; see also SI).

The specific feature-sets that emerged as those that shape the connectome in each of the three systems we studied partially overlap but are not identical (Fig. 4b). This is partly due to the available experimental data information (e.g., information on the dendritic tree length was only available for the OB data), but, potentially, also due to differences in the network structure in these circuits. For example, the OB connectome in the fish is highly reciprocal, whereas the cortical data from the mouse is not. Moreover, we found cell-type-specific connectivity to be the most important feature for accurately predicting the connectome of *C. elegans*, while reciprocity and preferential attachment within glomeruli were the dominant features in the OB ensemble. The connectivity in the mouse visual cortex connectome was highly directional (i.e., spatially-modulated) and asymmetric, and was accurately described by a model that includes a quadratic relation between a neuron’s spatial coordinates and its incoming and outgoing synapses (Methods).

We further validated our models by comparing their predictions of synaptic connectivity and the experimental estimates of the synaptic weights of the reconstructed connections. As our models relied only on the existence or absence of synaptic connections (i.e., binary connectivity matrices), we asked how well did the predictions of our models for the existence of a connection agree with the empirical estimations of individual synaptic strength. As shown in Fig. 4d, in all three species, the probability that our models assigned to a synapse strongly correlated with the measured weights.

Given the ability of our models to accurately predict individual synapses, we proceeded to test their accuracy in predicting the connectivity patterns of individual neurons and those of small subnetworks.

### Models accurately predict neuronal and circuit level properties

We evaluated the ability of our models to go beyond individual synapses, and predict the indegree and out-degree profiles of neurons in all three connectomes. We generated 500 synthetic connectomes from each of the selected models for each species, and compared their indegree and outdegree distributions to the empirical distributions in the experimental data. We found high agreement between the model predictions and the empirical degree distributions for all three connectomes, with empirical degree counts falling within the 90% confidence interval of the model. There was some discrepancy in the case of low-degree neurons in the OB connectome, which were predicted to be less common than the measured ones (Fig. 5a,b). Our models also predicted the existence of high-degree nodes or “hub neurons”. For example, according to the model, the probability of finding OB neurons whose indegree was equal or larger than the 90% percentile of indegrees in the data was between 4.0%-6.7% (compared with 0% for the ER model). Importantly, this prediction of the models did not rely on any specific feature for “hubbiness”; rather, it emerged from the more local and simple features of the model.

**Fig. 5.**
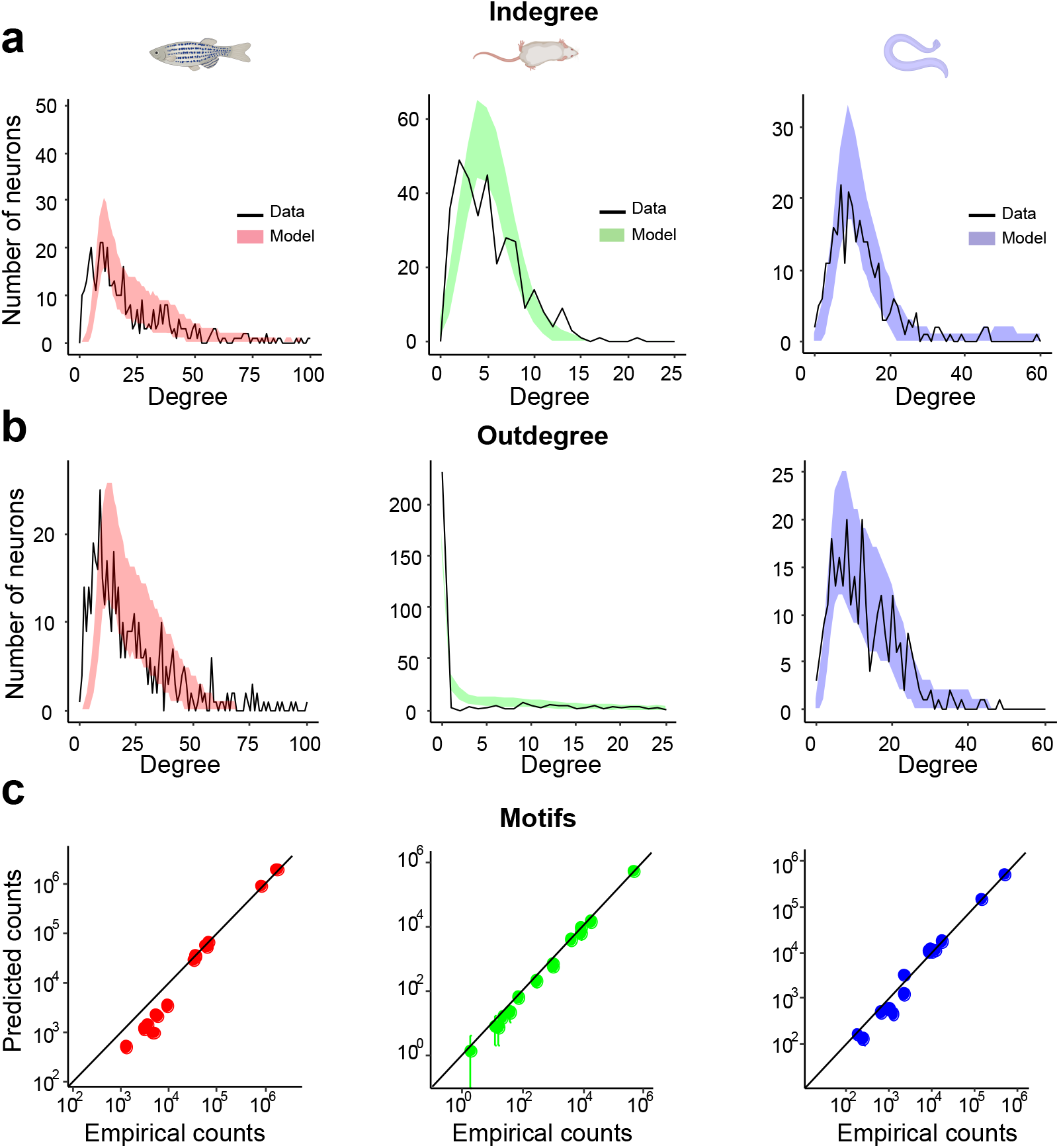
Connectome models accurately predict neuronal and circuit level properties in zebrafish, mouse, and *C. elegans.* **(a)** The Indegree distributions predicted by the selected models for all three connectomes. Black line - empirical indegree/outdegree distributions. Colored histograms - the 5th and 95th percentile computed from 500 samples from each model. **(b)** Same as in (a) but for the outdegree distribution. **(c)** Predictions of triplet network motifs. Error bars correspond to the 5% and 95% quantiles across 500 samples from each model.

We then evaluated the ability of our models to predict properties beyond those of single neurons. We first computed the distribution of all the connectivity patterns among triplets of neurons (“network motifs”) for each of the connectomes and models, as the frequency of these motifs has been suggested to be important for the kind of computations biological networks perform (10). We found that our models accurately predict the frequency of small network motifs, despite none of the models being constrained to reproduce them directly (the median normalized difference between the motif counts predicted by the model and the empirical counts was 18%; see Fig. 5c and Methods). We also estimated the distribution of the shortest-path distances between neurons, which describes how quickly information can flow between different parts of the network. We found high agreement between the shortest-path predictions of the models and the empirical data (Supplementary Fig. S8), including an accurate prediction of the number of pairs without a directed synaptic path between them (i.e., infinite shortest path distance). Thus, relying on a small set of physical and biological features of single neurons and pairwise relations, our models replicate, across species, a wide range of network properties – from distributions of indegrees and outdegrees, through the shortest-distance between neurons, to triplet motifs. Given this degree of accuracy, we next asked whether our models capture not just the structure of real connectomes but also the computation carried out by real circuits.

### Connectomes synthesized by the model reproduce the computation of real circuits

The OB circuit in the zebrafish is known to decorrelate the population activity representing odors and to normalize activity across the population (42–44). A two-step model based on the measured connectivity reproduces the “whitening” computation (34): the first step uses the activity of the MC neurons after stimulus presentation (time *t*_1_) to simulate the activity of the inhibitory population and the connectivity from MC→IN; the second step uses the predicted response of the inhibitory neurons, after thresholding and renormalization, and the connectivity from the IN→MC to simulate the activity of the excitatory population at the next time point, *t*_2_ (Fig. 6a). As the decorrelation between the predicted responses of the excitatory cells to different stimuli according to this model matches the experimentally measured one, we asked here whether networks synthesized by our generative models for the OB connectome would perform this whitening computation. In addition to focusing on our model of choice, we also asked how would networks synthesized by connectome models based on different features perform in response to olfactory stimuli.

**Fig. 6.**
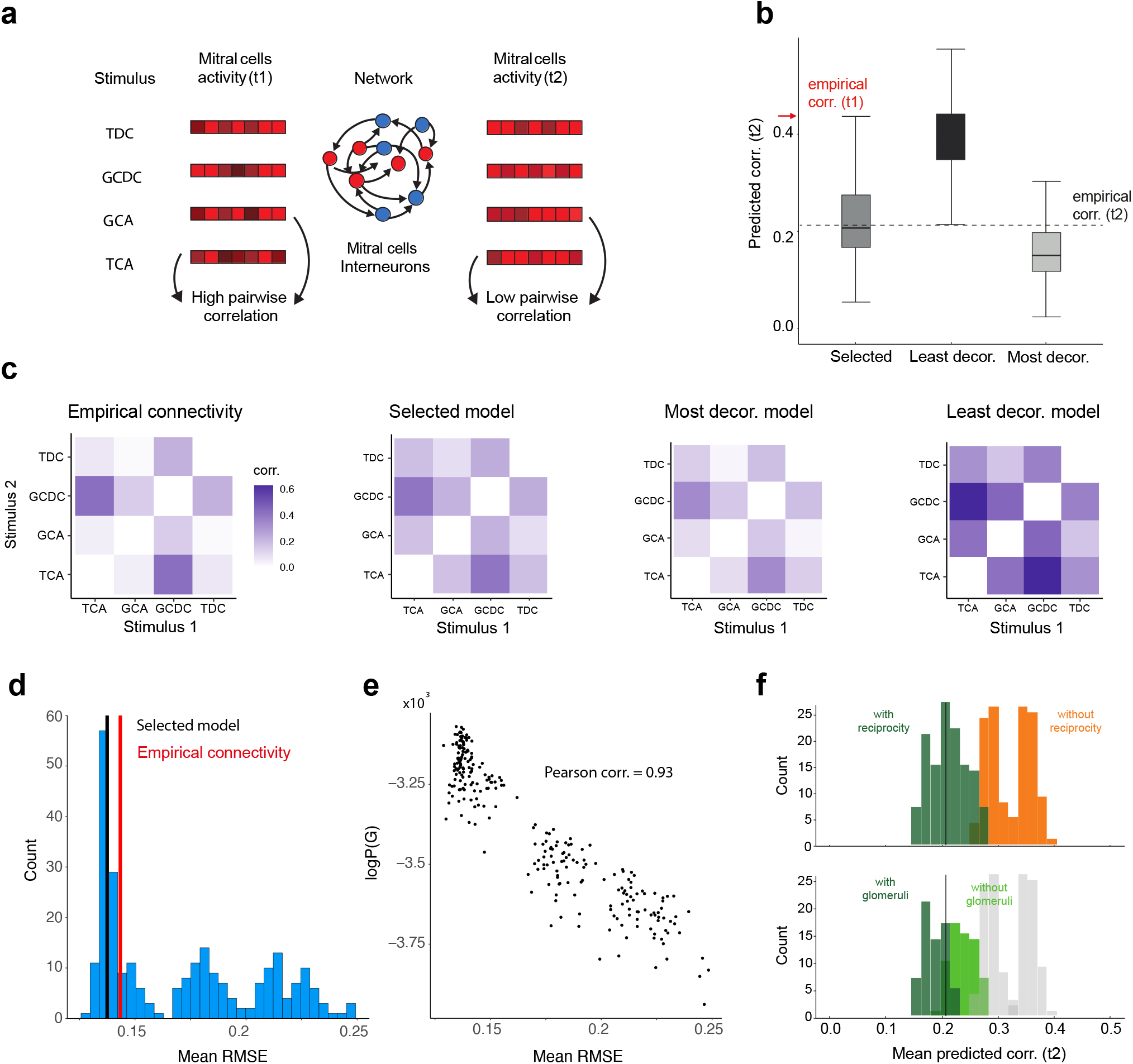
Connectome model generates neural circuits that recapitulate the computation performed by the olfactory bulb in zebrafish. **(a)** Simple model of the transformations of MC activity patterns by feedback inhibition (following (34): Input activity patterns (MC activity at *t*_1_)were multiplied by the feed-forward connectivity matrix MC→IN, then normalized, with the result thresholded at 2.8. The resultant IN activity patterns were multiplied by the feedback connectivity matrix IN→MC, yielding odorspecific patterns of feedback inhibition, which were subtracted from the MC activation patterns. Scaling factors and thresholds were adjusted such that the effects on the mean activity were small (adapted from Extended Data Figure 5 in (34)). **(b)** The distribution of the mean correlation at *t*_2_ (averaged across all six odor pairs) of 500 samples from the selected model (left), as well as the least decorrelating model (middle) and most decorrelating model (right). **(c)** The correlation matrix between four different odors, predicted by the simulation using (from left to right) the observed connectome, the selected model, and the most decorrelating and least decorrelating models. **(d)** The mean RMSE between the empirical pairwise correlations, computed from calcium traces, and the ones predicted by the model, for all models in the ensemble. For each model, the RMSE values were averaged over 500 samples from that model. Black vertical line shows the mean RMSE of the selected model; red vertical line corresponds to the RMSE computed using the empirical connectivity data. **(e)** The average RMSE (same as in (d)) vs. the log-likelihood of the data for each of the models in the ensemble. **(f)** The mean correlation values at *t*_2_ for all the models in the OB ensemble. Top - models are colored by whether they control for reciprocity (green) or not (orange). Bottom - models that control for reciprocity are colored by whether they also control for preferential attachment within glomeruli (dark green) or not (light green). Black vertical line corresponds to the empirical correlation at *t*_2_.

For each of the models in the above ensemble of 256 OB models (Fig. 3), we sampled 500 connectomes, and then simulated each network’s response to stimuli. For each network, we computed the correlation between the activity of the MC neurons at time *t*_2_ over different stimuli. Our results show that the models varied substantially in their ability to reproduce the observed decorrelation of MC neurons: Networks synthesized by some models showed no decorrelation, whereas others decorrelated more than what was observed empirically. We found, however, that the OB model we used to accurately predict the connectome was on par with the empirical connectome in its ability to decorrelate stimulus representations (Fig. 6b). Moreover, the networks synthesized by this model replicated the empirical decorrelation pattern – not only on average but also at the level of individual odor pairs (Fig. 6c-d, Supplementary Fig. S9). We further found that the ability of each of the models to replicate the measured connectome, as measured by the model’s likelihood, strongly correlated (Pearson correlation of 0.03, p-value < 10^−^ 6) with the similarity of the model’s responses to the empirical decorrelation of the network’s response to stimuli (Fig. 6e). Thus, the more accurate our structural model was, the better it replicated the computation that the circuit performed.

Comparing the performances of the different models also allowed us to identify the role of individual structural features in shaping the function of the network. We found that models that included the reciprocity in connectivity between neurons were very different than those that did not in their ability to perform the whitening computation (Fig. 6f, top). Interestingly, this was also the case for the likelihood values of the models. Out of the models that include reciprocity, the ones that also include preferential attachment within glomeruli showed stronger decorrelation than those that did not (Fig. 6f, bottom). Therefore, our results reflect the importance of these two structural properties in the context of the whitening computation.

### Model inaccuracies reveal new design features of the connectome

The selected feature-based models we presented offer a highly accurate “working draft” of the design principles of the connectomes we studied, yet they can probably be improved, up to the inherent biophysical noise in neuronal wiring and biological diversity across individuals. Improving such models could be achieved by assuming or guessing which new features one might add to the model. However, a particular strength of our framework is that the discrepancies between the model and the data must rely on “orthogonal” or independent features (see below), and this can be used to generate new hypotheses of additional design mechanisms. As an example, we focus here on the connectivity between 69 ventral cord motor neurons in *C. elegans,* which are connected by 352 chemical synapses. The left panel in Fig. 7a shows that when this connectivity matrix is ordered by the names of the neurons, distinct diagonal connectivity patterns within specific sub-matrices are apparent. The model we presented for the connectome of the whole worm in Fig. 3 captures some of these structures (namely, the blocks within the matrix that show a higher synaptic probability on the diagonal of each block), but fails to differentiate between blocks that contain synapses and blocks that do not (Fig. 7a, middle).

**Fig. 7.**
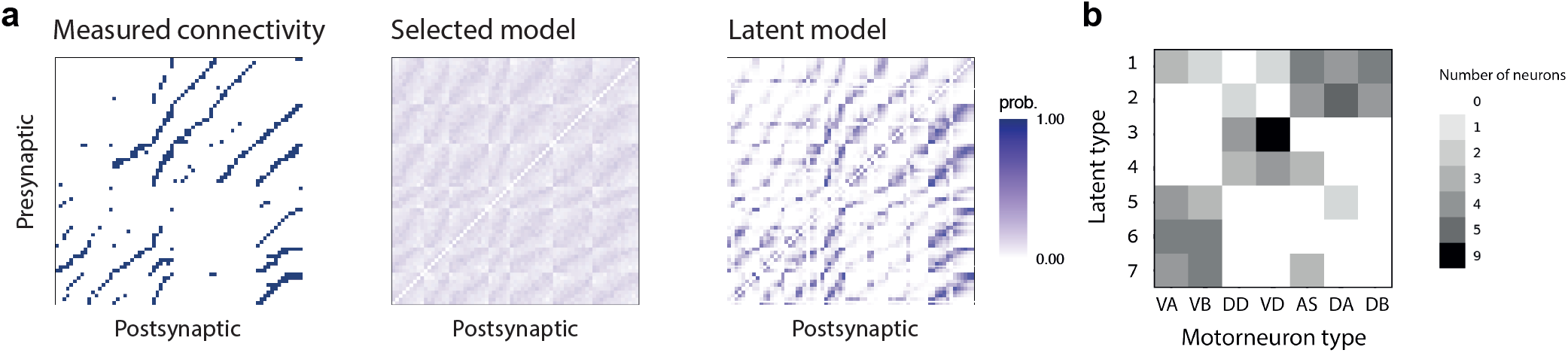
Inaccuracies of the models can be used to reveal missing design features of the connectome. **(a)** Connectivity between 69 ventral cord motor neurons in the worm: empirical connectivity data (left), synaptic probabilities according to the selected model (middle), and synaptic probabilities according to a distance+latent-types model of the circuit (right). **(b)** 2D histogram of motor neuron classes and inferred latent classes. Each entry in the matrix corresponds to the number of neurons of a given motorneuron type (x-axis) that were assigned by the optimization procedure to the same latent type (y-axis). Darker colors correspond to a larger number of neurons.

Because our model is a maximum entropy one, it gives the unique minimal structured network given the features it uses, and does not assume or imply other structural features. Thus, any discrepancies between a model based on some set of features and the data means that the model is missing some features. We, therefore, asked whether the observed structure of this part of the connectome corresponds to sub-types among the ventral cord motor neurons. We inferred such latent classes directly from the connectivity data by maximizing the likelihood of a model of the connectome, by iteratively adding features that correspond to *k* = 2,…, 10 potential sub-classes of neurons, while using the same distance-decay parameter as the selected model (see Methods). We found a highly accurate model for the motor neurons’ connectivity (Fig. 7a, right), with a maximum AUROC of 0.92 for *k* = 7 sub-types. Our latent classes could correspond to different cell types, morphological structures, as well as other synaptic specificity mechanisms. To identify the biological nature of the inferred latent classes, we inspected the specific neurons that were assigned to each latent class (Fig. 7b). It was particularly satisfying to find a clear correspondence between the latent classes and a subdivision of the neurons of the circuit according to their ventral/dorsal position, as well as their class (A/B/D class ventral cord motor neurons). Thus, our modeling approach allowed us to identify specific parts of the connectome where design features were missing from the model, and then to predict which biological features might be missing – which in this particular case, we could actually corroborate.

## Discussion

We present a family of generative statistical models for connectomes of three different neural circuits in three different species. In all cases, a small set of local biological and physical features were enough to accurately predict a range of structural measures of the real connectomes at the level of synaptic, neuronal, sub-circuit, and whole-network properties. In one case, measurements of the circuit before its structural reconstruction allowed us to show that the connectomes that the model generates replicate the computation performed by the real circuit.

Our results reflect that a surprisingly small set of features shapes much of the architectural design of connectomes. Moreover, given the stochastic nature of individual connectomes, our models may be even better than our results seem to suggest, since their accuracy is bounded by the biological variability between individuals. Importantly, the framework we present can be naturally extended to learning models of individual and populations of animals, once such data becomes available.

It is clear that at least part of the inaccuracies of the current models are due to the models missing some biological, regulatory, or physical features that play important roles in shaping connectomes. Measurements of additional morphological features or the delineation of more cell-types will probably result in even more accurate models. We note, however, that the fact that our models are generative allowed us to use them to identify where our models fail, and find latent features to add to the model and predict their biological nature – as we have shown here for the motor neurons of the worm.

We emphasize that while the features we used accurately describe the observed structure, this does not mean we have identified the true underlying mechanisms that control networks’ development and structure. Specifically, different subsets of features can define very similar probability distributions (as we have seen), in the same sense that different sets of vectors can span the same space. Thus, our modeling approach identifies potential sets of features that shape the connectome, but direct experimental characterization is needed to conclusively validate them or identify their alternatives.

Our modeling approach could be used to predict how experimental perturbations to synaptic formation mechanisms or to morphological properties of cells might affect large-scale organization and functional properties of the circuit (45). Probabilistic network models could also be used to predict which parameters are “stiff” and which are “sloppy” (46, 47), corresponding to different degrees of biological regulation, and how perturbations to one mechanism may be compensated by other mechanisms (48).

The networks we studied here are “snapshots” of the connectivity in one particular time point, and a natural extension would be to construct generative statistical models for the development of the connectome over time. While tracking the connectome of the same neural circuit over time is currently impossible (but see (5)), first steps have been made in analyzing connectomes of different individuals across development (49).

While we here focus on identifying design principles of the architecture of neural circuits, the ultimate goal is to identify the function these designs aim to achieve, which is often only partially known to external observers, and sometimes not at all. Extending our generative models beyond capturing the architectural design, to include computational goals for the circuits would be paramount for understanding the relations between the structure and function of neural networks – both biological and artificial ones. Ultimately, such models would allow the study and understanding of the structural basis of functional pathology in neural circuits and how to fix them.

## Acknowledgements

We thank the members of the Schneidman lab for comments, ideas, and discussions, and Andreas Tolias for sharing the cortical data with us, and advice. This work was supported by Simons Collaboration on the Global Brain grant 542997 (ES), Israel Science Foundation grant 137628 (ES), Israeli Council for Higher Education/Weizmann Data Science Research Center (ES), Swiss National Science Foundation 31003A_172925/1 (RF), the European Research Council (ERC) under the European Union’s Horizon 2020 research and innovation program (grant agreement 742576) (RF), Martin Kushner Schnur, and Mr. & Mrs. Lawrence Feis. ES is the incumbent of the Joseph and Bessie Feinberg Chair.

## Methods

### Feature-based maximum entropy models of connectomes

The maximum entropy framework defines the most random distribution *p*(*x*) that obeys a set of constraints of the form 〈*f_μ_*〉_*p*_ = 〈*f_μ_*〉_*data*_. The solution is given by solving the constrained optimization problem (50):

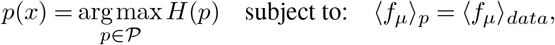

where *H*(*p*) is the Shannnon entropy *H*(*p*) = ∑_*x*_ *p*(*x*)log*p*(*x*) and 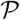 is the space of all the probability distributions over the relevant support. The optimal solution then takes the form of an exponential distribution:

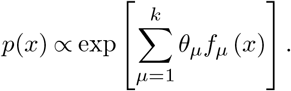

Exponential Random Graph Models (ERGMs) are a class of statistical network models for specifying a maximum entropy probability distribution over a set of random graphs of *N* nodes. An ERGM specifies the probability of a given network as a function of a set of structural features of the network, weighted by the model parameters. Importantly, for a given set of structural features, there is a one-to-one mapping from the set of parameter vectors {*θ*|*θ* ∈ ℝ^*k*^} to the set of realizable expectation values of the features {〈*f*_1_〉,…,〈*f_k_*〉}. Learning the parameters of a model, given a set of structural features and their values for a given graph, means finding the vector of parameters that would reproduce the expectation values of the observed features (50).

### Mathematical definitions of structural features

The structural features used to construct the various models in this paper (see main text) are described by the following mathematical formulas (where *G* is a binary *N* x *N* matrix with zeros on the main diagonal):

- Number of synapses: ∑_*i*≠*j*_ *G_ij_*
- Number of reciprocal synapses: ∑_*i*>*j*_ *G_ij_G_ji_*
- Distance between connected neurons: ∑_*i*≠*j*_ *d*(*i,j*)· *G_ij_*, where *d*(*i,j*) is the distance between neuron *i* and neuron *j*.
- Number of synapses between neurons of type *A* and type 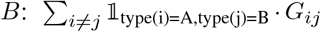: each combination of cell types has a different model parameter (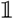 is the indicator function).
- Effect of single-cell properties on a neuron’s tendency to receive incoming synapses: ∑_*i*≠*j*_ *h*(*j*) · *G_ij_*, where *h*(*j*) is a property of the *j*-th neuron. The properties we considered are the spatial coordinates of the neurons, the size of their dendritic trees, and their time of birth.
- Effect of single-cell properties on a neuron’s tendency to form outgoing synapses - same equation as the previous one, with *h*(*i*) instead of *h*(*j*).
- Number of synapses whose pre-synaptic and post-synaptic neurons belong to the same glomerulus: 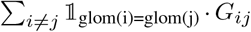. A similar feature was used to assigned the microdomain of pre- and post-synaptic neurons.

### Connectomics data preprocessing

The full connectome of the olfactory bulb of the zebrafish, previously reported in (34), consists of 1003 cells. Of these, 467 neurons were used in this study: the 208 MCs for which activity data were available and the 259 INs connected to these MCs. After removing disconnected cells and cells whose dendritic tree size was not reconstructed, 446 cells remained. Rows and columns were ordered as in (34). The connectome of *C.elegans* was based on data from (40). Our analysis focused on the connectivity of chemical synapses (and not electric gap junctions) in the hermaphrodite. For the purposed of the analysis, we omitted from the original data, which consisted of 300 neurons, the pharyngeal neurons, focusing on 279 sensory neurons, inter-neurons and motor neurons for which we had additional neuron-level data (e.g., spatial position; see main text). Rows and columns were ordered as in (40). 3D cell positions were obtained from (51). The connectome of the mouse was based on a publicly available dataset, previously reported in (39). Rows and columns were ordered by optimal leaf ordering for hierarchical clustering. Autapses were excluded in all 3 datasets. For additional details, see the accompanying GitHub repository: https://github.com/adamhaber/ergm_for_connectomics

### Fitting and sampling

All models were fitted using the ERGM R library, version 4.1.2 (52). Finding the maximumlikelihood estimator (MLE) of a model that assumes independence between different edges, given the covariates of the model, is equivalent to solving a logistic-regression (53). Similarly, such models are amenable to exact sampling, since synapses are sampled independently according to the marginal synaptic probabilities, which can be computed analytically. Finding the MLE of a model that assumes edges are dependent (e.g., all models that control for the observed level of reciprocity) is done via Monte Carlo maximum likelihood estimation (MCMLE), a likelihood-based approach that relies on Markov Chain Monte Carlo (MCMC) sampling for estimating the parameters of the model. Similarly, sampling is done via the Metropolis-Hastings algorithm, as described in (54).

### Computing the matrix of synaptic probabilities

For independent models, the matrix of synaptic probabilities can be computed analytically, as the predicted probabilities according to the logistic regression model that is used to infer the MLE (53). For dependent models (e.g., models with a reciprocity term), we used MCMC to sample 500 networks, using 10,000,000 burn-in iterations (number of proposals before any sampling is done) and an interval of 1,000,000 proposals between sampled networks. The connectivity matrices of the sampled networks were averaged to obtain the empirical matrix of synaptic probabilities.

### Cross-validation using subnetworks

For each connectome, we randomly assigned each neuron to one of two equally sized groups (of size 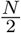, where *N* is the total number of neurons) - a train subnetwork and a test subnetwork. We fitted the model to the train network, and used the inferred MLE to compute the log-likelihood of the test network. We note that this required us to re-evaluate the normalizing constant, as the exogenous covariates (such as cell types) of the test network were generally different from those of the train network. We conducted 10-fold cross-validations for each model, for each dataset.

### Inferring latent motor neuron classes

To infer latent motor neuron classes, we employed a simple greedy optimization procedure. We began by randomly assigning each of the 69 motor neurons to one of *k* latent classes (we considered values of *k* between 2 and 10). A single iteration of the optimization procedure consisted of randomly choosing a single neuron and finding its latent class assignment that would maximally increase the log-likelihood of the data, while keeping the class assignment of all other 68 neurons fixed. The model’s performance was evaluated using the latent classes obtained after 10,000 optimization steps.

## References

1. Louis K Scheffer, C Shan Xu, Michal Januszewski, et al. A connectome and analysis of the adult Drosophila central brain. eLife, 9(6):e57443, 2020.

2. Alexander Shapson-Coe, Michał Januszewski, Daniel R. Berger, Art Pope, Yuelong Wu, Tim Blakely, Richard L. Schalek, Peter Li, Shuohong Wang, Jeremy Maitin-Shepard, Neha Karlupia, Sven Dorkenwald, Evelina Sjostedt, Laramie Leavitt, Dongil Lee, Luke Bailey, Angerica Fitzmaurice, Rohin Kar, Benjamin Field, Hank Wu, Julian Wagner-Carena, David Aley, Joanna Lau, Zudi Lin, Donglai Wei, Hanspeter Pfister, Adi Peleg, Viren Jain, and Jeff W. Lichtman. A connectomic study of a petascale fragment of human cerebral cortex. biorxiv, 2021. doi: 10.1101/2021.05.29.446289.

3. Alessandro Motta, Manuel Berning, Kevin M. Boergens, Benedikt Staffler, Marcel Beining, Sahil Loomba, Philipp Hennig, Heiko Wissler, and Moritz Helmstaedter. Dense connectomic reconstruction in layer 4 of the somatosensory cortex. Science, 366(6469):eaay3134, 2019.

4. Sahil Loomba, Jakob Straehle, Vijayan Gangadharan, Natalie Heike, Abdelrahman Khalifa, Alessandro Motta, Niansheng Ju, Meike Sievers, Jens Gempt, Hanno S. Meyer, and Moritz Helmstaedter. Connectomic comparison of mouse and human cortex. Science, 377(6602): eabo0924, 2022.

5. Xiaoyin Chen, Yu-Chi Sun, Huiqing Zhan, Justus M Kebschull, Stephan Fischer, Katherine Matho, Z Josh Huang, Jesse Gillis, and Anthony M Zador. High-throughput mapping of long-range neuronal projection using in situ sequencing. Cell, 179(3):772–786, 2019.

6. Nicholas L Turner, Thomas Macrina, J Alexander Bae, Runzhe Yang, Alyssa M Wilson, Casey Schneider-Mizell, Kisuk Lee, Ran Lu, Jingpeng Wu, Agnes L Bodor, et al. Reconstruction of neocortex: Organelles, compartments, cells, circuits, and activity. Cell, 185: 1082–1100, 2022.

7. Larry F. Abbott, Davi D. Bock, Edward M. Callaway, Winfried Denk, Catherine Dulac, Adrienne L. Fairhall, Ila Fiete, Kristen M. Harris, Moritz Helmstaedter, Viren Jain, Narayanan Kasthuri, Yann LeCun, Jeff W. Lichtman, Peter B. Littlewood, Liqun Luo, John H. R. Maunsell, R. Clay Reid, Bruce R. Rosen, Gerald M. Rubin, Terrence J. Sejnowski, H. Sebastian Seung, Karel Svoboda, David W. Tank, Doris Tsao, and David C. Van Essen. The Mind of a Mouse. Cell, 182(6):1372–1376, 2020.

8. Anastasia A. Makarova, Alexey A. Polilov, and Dmitri B. Chklovskii. Small brains for big science. Current Opinion in Neurobiology, 71:77–83, 2021.

9. Sen Song, Per Jesper Sjöström, Markus Reigl, Sacha Nelson, and Dmitri B Chklovskii. Highly nonrandom features of synaptic connectivity in local cortical circuits. PLoS biology, 3(3):e68, 2005.

10. R Milo, S Shen-Orr, S Itzkovitz, N Kashtan, D Chklovskii, and U Alon. Network motifs: simple building blocks of complex networks. Science, 298(5594):824–827, 2002.

11. Brad K Hulse, Hannah Haberkern, Romain Franconville, Daniel Turner-Evans, Shin-ya Takemura, Tanya Wolff, Marcella Noorman, Marisa Dreher, Chuntao Dan, Ruchi Parekh, Ann M Hermundstad, Gerald M Rubin, and Vivek Jayaraman. A connectome of the *Drosophila* central complex reveals network motifs suitable for flexible navigation and context-dependent action selection. eLife, 10:e66039, oct 2021.

12. Jonathan Green, Atsuko Adachi, Kunal K. Shah, Jonathan D. Hirokawa, Pablo S. Magani, and Gaby Maimon. A neural circuit architecture for angular integration in drosophila. Nature, 546:101–106, oct 2017.

13. Moritz Helmstaedter, Kevin L. Briggman, Srinivas C. Turaga, Viren Jain, H. Sebastian Seung, and Winfried Denk. Connectomic reconstruction of the inner plexiform layer in the mouse retina. Nature, 500(7461):168–174, 2013.

14. MICrONS Consortium, J Alexander Bae, Mahaly Baptiste, Agnes L Bodor, Derrick Brittain, JoAnn Buchanan, Daniel J Bumbarger, Manuel A Castro, Brendan Celii, Erick Cobos, et al. Functional connectomics spanning multiple areas of mouse visual cortex. bioRxiv, 2021. doi: 10.1101/2021.07.28.454025.

15. Narayanan Kasthuri, Kenneth Jeffrey Hayworth, Daniel Raimund Berger, Richard Lee Schalek, José Angel Conchello, Seymour Knowles-Barley, Dongil Lee, Amelio Vázquez-Reina, Verena Kaynig, Thouis Raymond Jones, et al. Saturated reconstruction of a volume of neocortex. Cell, 162(3):648–661, 2015.

16. Casey M Schneider-Mizell, Agnes Bodor, Derrick Brittain, JoAnn Buchanan, Daniel J. Bumbarger, Leila Elabbady, Daniel Kapner, Sam Kinn, Gayathri Mahalingam, Sharmishtaa Seshamani, Shelby Suckow, Marc Takeno, Russel Torres, Wenjing Yin, Sven Dorkenwald, J. Alexander Bae, Manuel A. Castro, Paul G. Fahey, Emmanouil Froudakis, Akhilesh Halageri, Zhen Jia, Chris Jordan, Nico Kemnitz, Kisuk Lee, Kai Li, Ran Lu, Thomas Macrina, Eric Mitchell, Shanka Subhra Mondal, Shang Mu, Barak Nehoran, Stelios Papadopoulos, Saumil Patel, Xaq Pitkow, Sergiy Popovych, William Silversmith, Fabian H. Sinz, Nicholas L. Turner, William Wong, Jingpeng Wu, Szi-chieh Yu, Jacob Reimer, Andreas S. Tolias, H Sebastian Seung, R Clay Reid, Forrest Collman, and Nuno Maçarico da Costa. Cell-type-specific inhibitory circuitry from a connectomic census of mouse visual cortex. bioRxiv, 2023. doi: 10.1101/2023.01.23.525290.

17. Lee Cossell, Maria Florencia Iacaruso, Dylan R. Muir, Rachael Houlton, Elie N. Sader, Ho Ko, Sonja B. Hofer, and Thomas D. Mrsic-Flogel. Functional organization of excitatory synaptic strength in primary visual cortex. Nature, 518(7539):399–403, 2015.

18. Yushu Chen, Xiaoyin Chen, Batuhan Baserdem, Huiqing Zhan, Yan Li, Martin B. Davis, Justus M. Kebschull, Anthony M. Zador, Alexei A. Koulakov, and Dinu F. Albeanu. High-throughput sequencing of single neuron projections reveals spatial organization in the olfactory cortex. Cell, 185(22):4117–4134, 2022.

19. Christopher A. Brittin, Steven J. Cook, David H. Hall, Scott W. Emmons, and Netta Cohen. A multi-scale brain map derived from whole-brain volumetric reconstructions. Nature, 591: 105–110, 2021.

20. István A Kovács, Dániel L Barabási, and Albert-László Barabási. Uncovering the genetic blueprint of the c. elegans nervous system. Proceedings of the National Academy of Sciences, 117(52):33570–33577, 2020.

21. Molly B Reilly, Cyril Cros, Erdem Varol, Eviatar Yemini, and Oliver Hobert. Unique homeobox codes delineate all the neuron classes of c. elegans. Nature, 584(7822):595–601, 2020.

22. Seth R Taylor, Gabriel Santpere, Alexis Weinreb, Alec Barrett, Molly B Reilly, Chuan Xu, Erdem Varol, Panos Oikonomou, Lori Glenwinkel, Rebecca McWhirter, et al. Molecular topography of an entire nervous system. Cell, 184(16):4329–4347, 2021.

23. Dániel L Barabási and Albert-László Barabási. A genetic model of the connectome. Neuron, 105(3):435–445, 2020.

24. Michael W Reimann, Anna-Lena Horlemann, Srikanth Ramaswamy, Eilif B Muller, and Henry Markram. Morphological diversity strongly constrains synaptic connectivity and plasticity. Cerebral Cortex, 27(9):4570–4585, 2017.

25. Daniel Udvary, Philipp Hart, Jakob H Macke, Hans-Christian Hege, Christiaan PJ de Kock, Bert Sakman, and Marcel Oberlaender. The impact of neuronal structure on cortical network architecture. bioRxiv, 2020. doi: 10.1101/2020.11.13.381087.

26. Joshua R Sanes and S Lawrence Zipursky. Synaptic specificity, recognition molecules, and assembly of neural circuits. Cell, 181(3):536–556, 2020.

27. Ed Bullmore and Olaf Sporns. The economy of brain network organization. Nature reviews neuroscience, 13(5):336–349, 2012.

28. M. E. J. Newman. The structure and function of complex networks. SIAM Review, 45(2): 167–256, 2003.

29. Albert-László Barabási and Réka Albert. Emergence of Scaling in Random Networks. Science, 286(5439):509 – 512, 1999.

30. Danielle S Bassett and Olaf Sporns. Network neuroscience. Nature neuroscience, 20(3): 353–364, 2017.

31. Richard F. Betzel and Danielle S. Bassett. Generative models for network neuroscience: prospects and promise. Journal of The Royal Society Interface, 14(136):20170623, November 2017.

32. Danielle Smith Bassett and ED Bullmore. Small-world brain networks. The neuroscientist, 12(6):512–523, 2006.

33. Richard F Betzel, Andrea Avena-Koenigsberger, Joaquín Goñi, Ye He, Marcel A De Reus, Alessandra Griffa, Petra E Vértes, Bratislav Mišic, Jean-Philippe Thiran, Patric Hagmann, et al. Generative models of the human connectome. Neuroimage, 124:1054–1064, 2016.

34. Adrian A. Wanner and Rainer W. Friedrich. Whitening of odor representations by the wiring diagram of the olfactory bulb. Nature Neuroscience, 23(3):433–442, March 2020.

35. Simone Holler, German Köstinger, Kevan AC Martin, Gregor FP Schuhknecht, and Ken J Stratford. Structure and function of a neocortical synapse. Nature, 591(7848):111–116, 2021.

36. Garry Robins, Pip Pattison, Yuval Kalish, and Dean Lusher. An introduction to exponential random graph (p*) models for social networks. Social networks, 29(2):173–191, 2007.

37. Edwin T Jaynes. Information theory and statistical mechanics. Physical review, 106(4):620, 1957.

38. Ori Maoz, Gašper Tkačik, Mohamad Saleh Esteki, Roozbeh Kiani, and Elad Schneidman. Learning probabilistic neural representations with randomly connected circuits. Proceedings of the National Academy of Sciences, 117(40):25066–25073, 2020.

39. Sven Dorkenwald, Nicholas L Turner, Thomas Macrina, Kisuk Lee, Ran Lu, Jingpeng Wu, Agnes L Bodor, Adam A Bleckert, Derrick Brittain, Nico Kemnitz, William M Silversmith, Dodam Ih, Jonathan Zung, Aleksandar Zlateski, Ignacio Tartavull, Szi-Chieh Yu, Sergiy Popovych, William Wong, Manuel Castro, Chris S Jordan, Alyssa M Wilson, Emmanouil Froudarakis, JoAnn Buchanan, Marc M Takeno, Russel Torres, Gayathri Mahalingam, Forrest Collman, Casey M Schneider-Mizell, Daniel J Bumbarger, Yang Li, Lynne Becker, Shelby Suckow, Jacob Reimer, Andreas S Tolias, Nuno Macarico da Costa, R Clay Reid, and H Sebastian Seung. Binary and analog variation of synapses between cortical pyramidal neurons. eLife, 11:e76120, nov 2022.

40. Steven J. Cook, Travis A. Jarrell, Christopher A. Brittin, Yi Wang, Adam E. Bloniarz, Maksim A. Yakovlev, Ken C. Q. Nguyen, Leo T.-H. Tang, Emily A. Bayer, Janet S. Duerr, Hannes E. Bülow, Oliver Hobert, David H. Hall, and Scott W. Emmons. Whole-animal connectomes of both Caenorhabditis elegans sexes. Nature, 571(7763):63–71, July 2019.

41. Emma K Towlson, Petra E Vértes, Sebastian E Ahnert, William R Schafer, and Edward T Bullmore. The rich club of the c. elegans neuronal connectome. Journal of Neuroscience, 33(15):6380–6387, 2013.

42. Rainer W Friedrich and Gilles Laurent. Dynamic optimization of odor representations by slow temporal patterning of mitral cell activity. Science, 291(5505):889–894, 2001.

43. Rainer W Friedrich and Martin T Wiechert. Neuronal circuits and computations: pattern decorrelation in the olfactory bulb. FEBS letters, 588(15):2504–2513, 2014.

44. Peixin Zhu, Thomas Frank, and Rainer W Friedrich. Equalization of odor representations by a network of electrically coupled inhibitory interneurons. Nature neuroscience, 16(11): 1678–1686, 2013.

45. Vladyslava Pechuk, Gal Goldman, Yehuda Salzberg, Aditi H. Chaubey, R. Aaron Bola, Jonathon R. Hoffman, Morgan L. Endreson, Renee M. Miller, Noah J. Reger, Douglas S. Portman, Denise M. Ferkey, Elad Schneidman, and Meital Oren-Suissa. Reprogramming the topology of the nociceptive circuit in c. elegans reshapes sexual behavior. Current Biology, 24(20):4372–4385, 2022.

46. Benjamin B Machta, Ricky Chachra, Mark K Transtrum, and James P Sethna. Parameter space compression underlies emergent theories and predictive models. Science, 342 (6158):604–607, 2013.

47. Ryan N Gutenkunst, Joshua J Waterfall, Fergal P Casey, Kevin S Brown, Christopher R Myers, and James P Sethna. Universally sloppy parameter sensitivities in systems biology models. PLoS computational biology, 3(10):e189, 2007.

48. E. Marder. Variability, compensation, and modulation in neurons and circuits. Proceedings of the National Academy of Sciences, 108:15542–15548, 2011.

49. Daniel Witvliet, Ben Mulcahy, James K. Mitchell, Yaron Meirovitch, Daniel R. Berger, Yuelong Wu, Yufang Liu, Wan Xian Koh, Rajeev Parvathala, Douglas Holmyard, Richard L. Schalek, Nir Shavit, Andrew D. Chisholm, Jeff W. Lichtman, Aravinthan D. T. Samuel, and Mei Zhen. Connectomes across development reveal principles of brain maturation. Nature, 596(7871):257–261, August 2021.

50. Martin J Wainwright, Michael I Jordan, et al. Graphical models, exponential families, and variational inference. Foundations and Trends in Machine Learning, 1(1–2):1–305, 2008.

51. Michael Skuhersky, Tailin Wu, Eviatar Yemini, Amin Nejatbakhsh, Edward Boyden, and Max Tegmark. Toward a more accurate 3d atlas of c. elegans neurons. BMC bioinformatics, 23 (1):1–18, 2022.

52. Pavel N Krivitsky, David R Hunter, Martina Morris, and Chad Klumb. ergm 4.0: new features and improvements. arXiv preprint arXiv:2106.04997, 2021.

53. Tom A B Snijders. Markov Chain Monte Carlo Estimation of Exponential Random Graph Models. page 40.

54. 54. Pavel N Krivitsky, David R Hunter, Martina Morris, and Chad Klumb. ergm 4: Computational improvements. arXiv preprint arXiv:2203.08198, 2022.

