## Supplemental Figures for "The structure and function of neural connectomes are shaped by a small number of design principles"

### Supplementary Figures

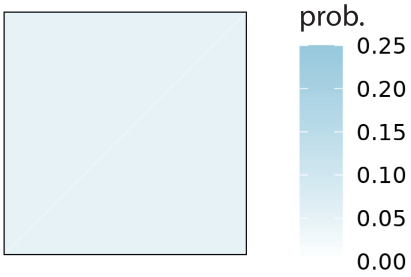


**Fig. S1. Synaptic predictions of the Erdős-Reńyi model.** Synaptic probabilities predicted by the Erdős-Reńyi model of the OB connectome, which only includes a number-of-synapses terms, adjusting for the overall sparsity of the network.

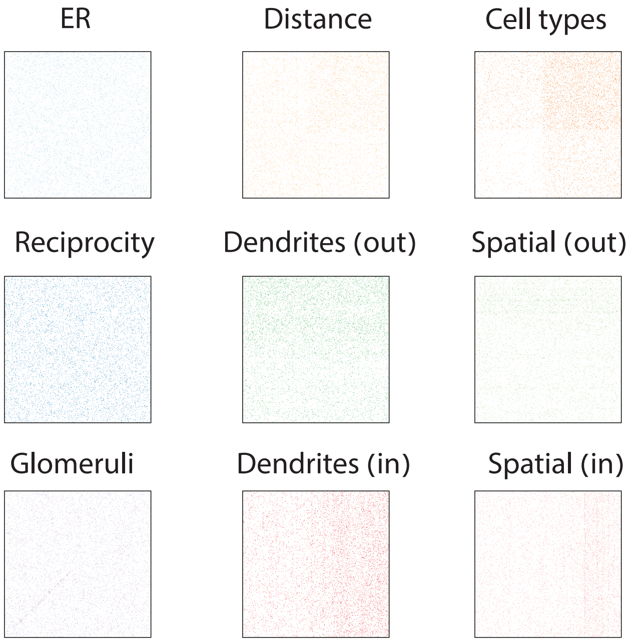


**Fig. S2. Individual samples of single feature models for the OB connectome.** Examples of individual samples drawn from the 8 models in Fig. 2b and the ER model from Supplementary Fig. S1 .

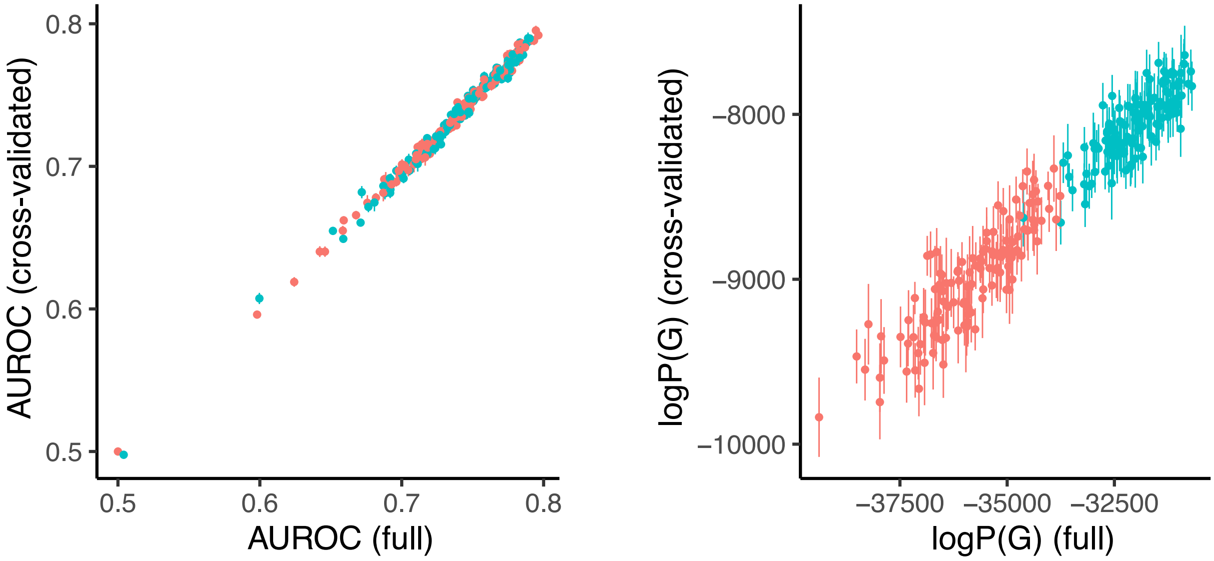


**Fig. S3. Cross-validated model scores are highly correlated with scores computed for the complete connectome. (a)** - Cross-validated AUROC (computed over 10 different train/test splits) vs. AUROC computed for the model trained on the entire connectome. Each point is a model in the OB models' ensemble. Cyan/red points correspond to models that include/do not include the reciprocity term, respectively. Error bars represent one standard deviation. **(b)** Same as in (a) but for models' log-likelihood scores.

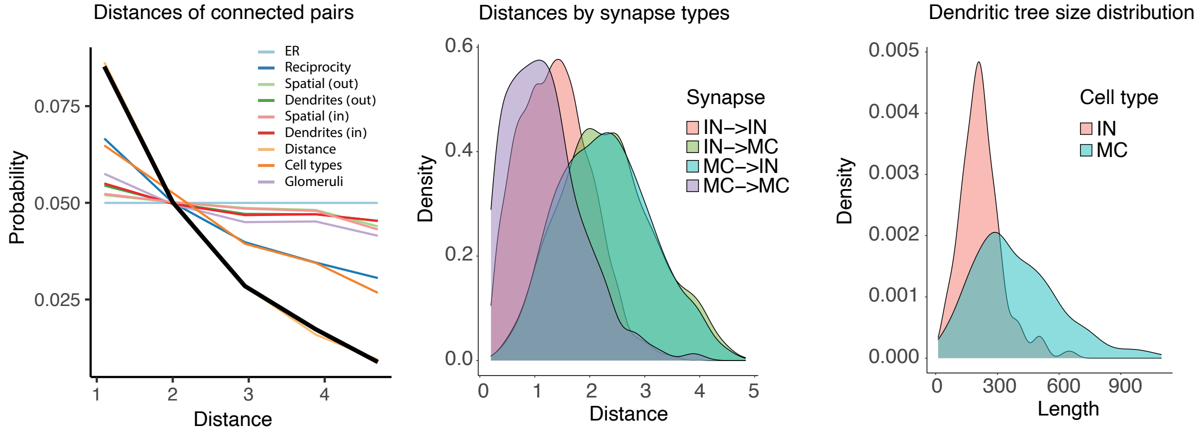


**Fig. S4. Correlations between different structural features in the OB connectome. (a)** The dependence of the probability of synaptic connections on the distance between cells, according to all 9 models based on single feature-sets (including ER). **(b)** Distributions of distances between pairs of neurons that have a synaptic connection between them, by the type of both the pre- and post-synaptic cells. **(c)** Size distributions of the dendritic tree, by cell type.

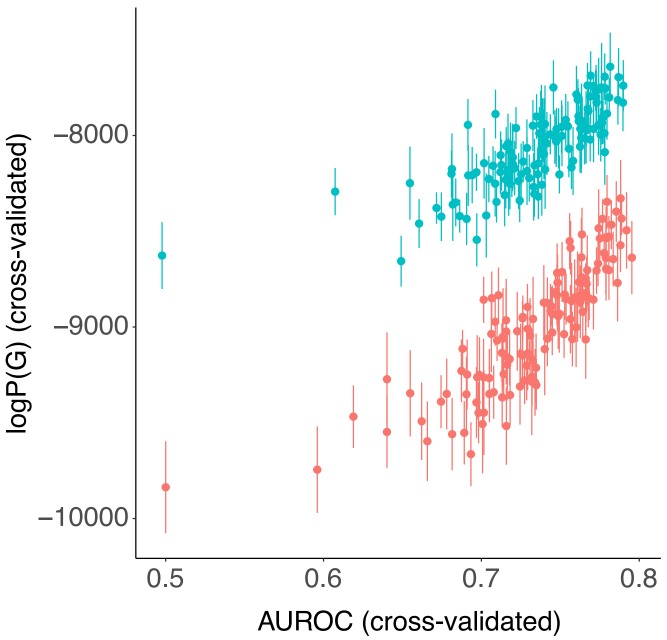


**Fig. S5. Correlation between the log-likelihood and AUROC scores.** Cross-validated AUROC scores vs. cross-validated log-likelihood scores. Each point is a model in the OB models ensemble. Cyan/red points correspond to models that include/do not include the reciprocity term, respectively. Error bars represent one standard deviation.

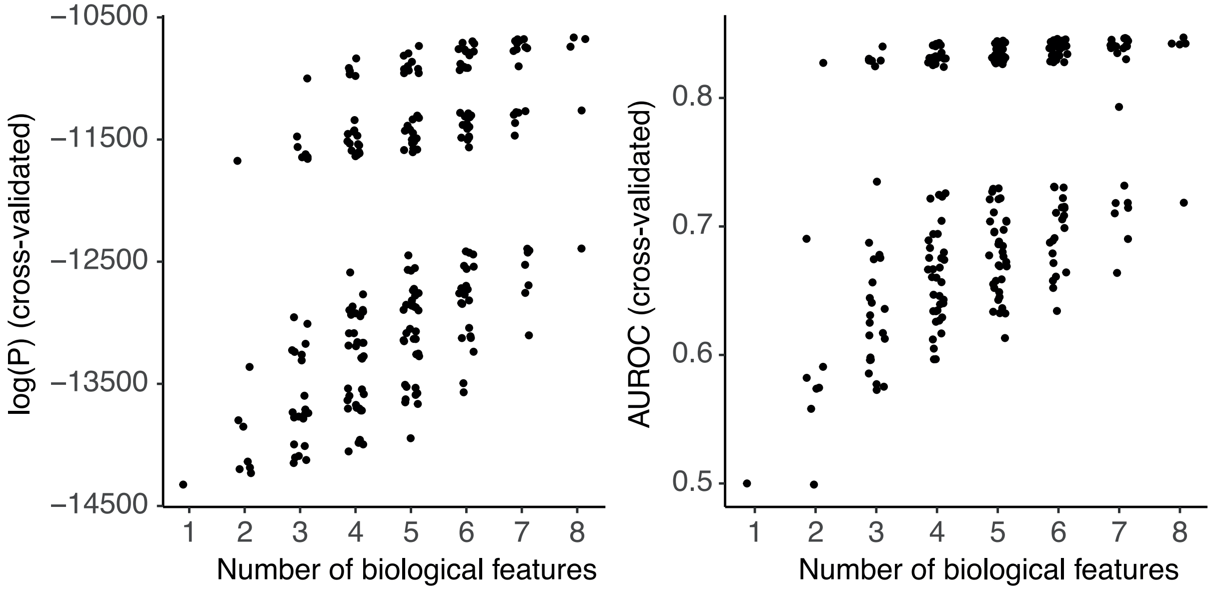


**Fig. S6. Performance of all Worm models.** The cross-validated log-likelihood (left) and AUROC scores (right) of all the models in the worm model ensemble.

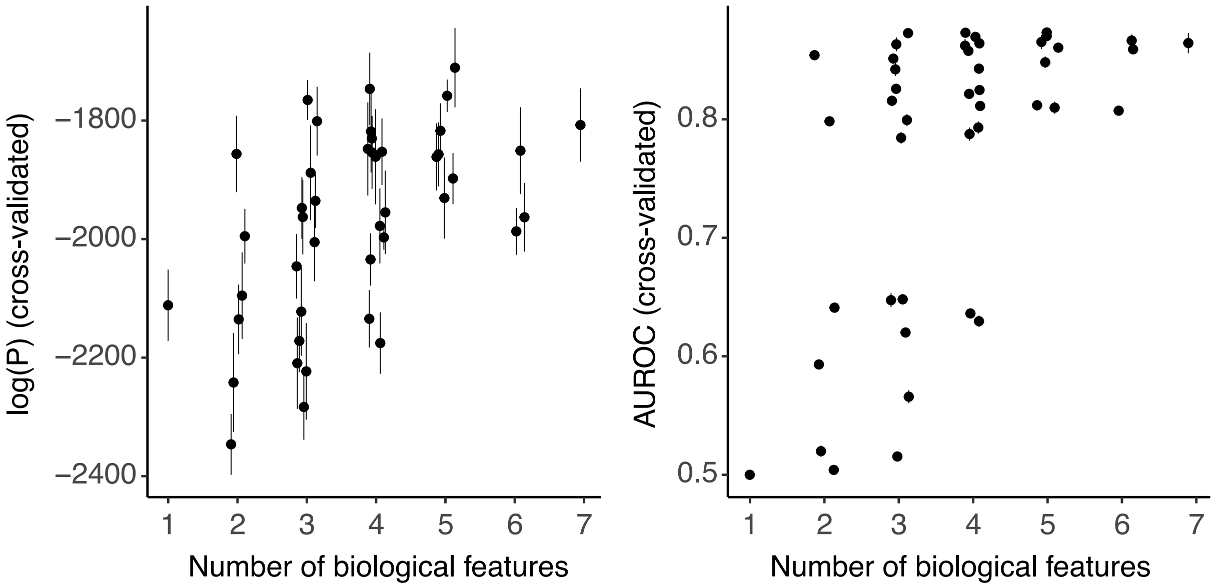


**Fig. S7. Performance of all Mouse models.** The cross-validated log-likelihood (left) and AUROC scores (right) of all the models in the mouse model ensemble.

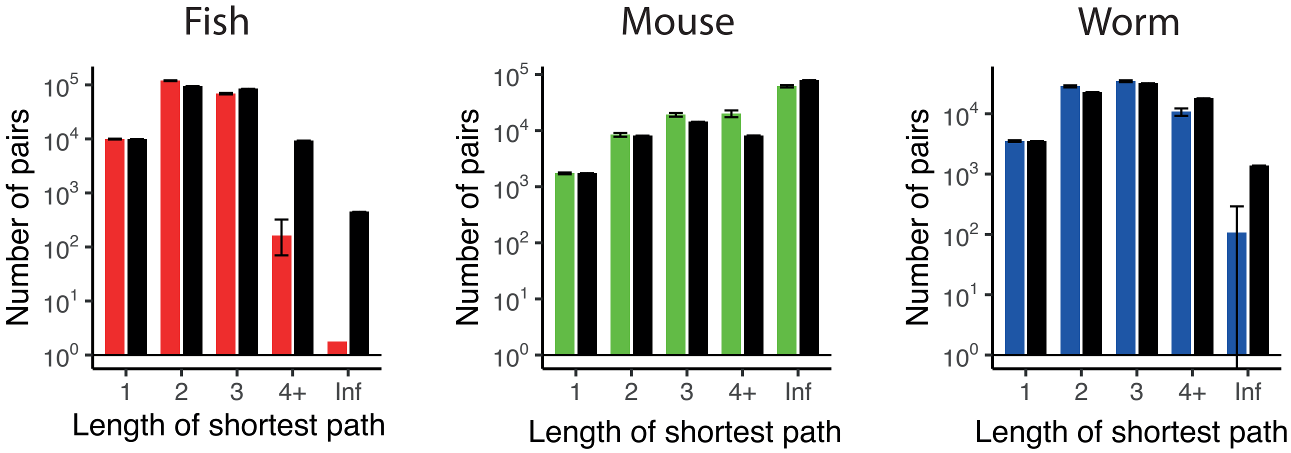


**Fig. S8. Distributions of shortest path between neuron pairs in all data sets and model predictions. (a)** Distribution of shortest path length between pairs of neurons. Inf means no path exists between the two neurons. Black - data; colored bars - predictions of the selected model for each dataset, computed from 500 samples. Error bars represent the 5% and 95% quantiles over the samples.

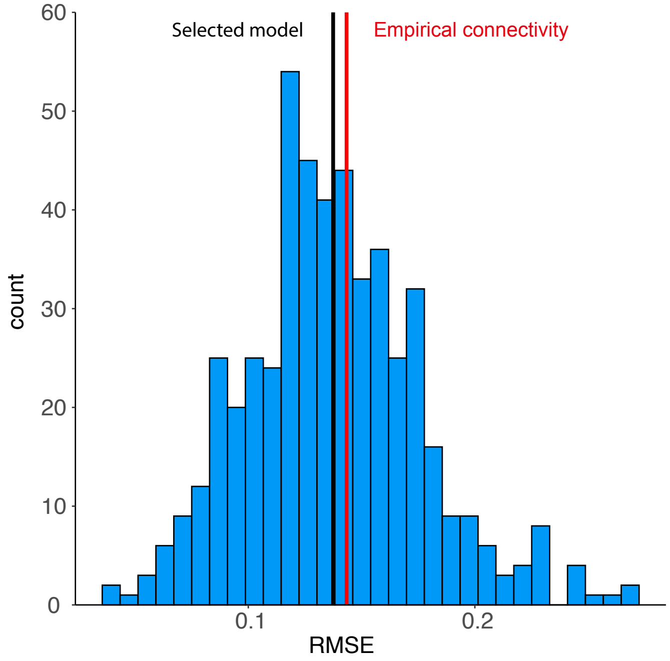


**Fig. S9. Accuracy of predicting the structure of pairwise correlations using samples from the selected OB model. (a)** Distribution of the RMSE between the empirical pairwise correlations at $t_2$ and those predicted by samples from the selected model (see main text), for 500 different samples. Black vertical line - average RMSE (same as in Fig.,6d); red - RMSE when using the empirical connectivity data to predict correlations at $t_2$.
